# Two Genomic Loci Control Three Eye Colors in the Domestic Pigeon (*Columba livia*)

**DOI:** 10.1101/2021.03.11.434326

**Authors:** Emily T. Maclary, Bridget Phillips, Ryan Wauer, Elena F. Boer, Rebecca Bruders, Tyler Gilvarry, Carson Holt, Mark Yandell, Michael D. Shapiro

## Abstract

The iris of the eye shows striking color variation across vertebrate species, and may play important roles in crypsis and communication. The domestic pigeon (*Columba livia)* has three common iris colors, orange, pearl (white), and bull (dark brown), segregating in a single species, thereby providing a unique opportunity to identify the genetic basis of iris coloration. We used comparative genomics and genetic mapping in laboratory crosses to identify two candidate genes that control variation in iris color in domestic pigeons. We identified a nonsense mutation in the solute carrier *SLC2A11B* that is shared among all pigeons with pearl eye color, and a locus associated with bull eye color that includes *EDNRB2*, a gene involved in neural crest migration and pigment development. However, bull eye is likely controlled by a heterogeneous collection of alleles across pigeon breeds. We also found that the *EDNRB2* region is associated with regionalized plumage depigmentation (piebalding). Our results establish a genetic link between iris and plumage color, two traits that were long known by pigeon breeders to co-occur, and demonstrate the importance of gene duplicates in establishing possibilities and constraints in the evolution of color and color pattern among vertebrates.

## INTRODUCTION

A wide variety of genetic and developmental mechanisms influence diversity in pigment type and patterning in the vertebrate epidermis, including epidermal appendages such as hair and feathers (Hoekstra 2006; Kelsh et al. 2008; Hubbard et al. 2010; Kaelin and Barsh 2013; Domyan et al. 2014; Parichy and Spiewak 2014; Bruders et al. 2020; Inaba and Chuong 2020). Pigments are also deposited in non-epidermal tissues in vertebrates, including the iris of the eye. Among vertebrate species, iris color varies widely. Some species have conspicuously colored bright yellow, red, or white irises, while others have dark irises. Iris coloration may be an adaptive trait that, like epidermal coloration, plays roles in crypsis and communication. For example, iris color is correlated with habitat in mantellid frogs, with arboreal species more likely to have bright eyes (Amat et al. 2013). Bright irises probably evolved multiple times in arboreal mantellid species, indicating that this trait might be adaptive. There is evidence that iris color may be adaptive in birds as well. In owls, dark iris color likely coevolved with nocturnal behavior (Passarotto et al. 2018), while the bright white irises of jackdaws communicate that nesting sites are occupied (Davidson et al. 2014; Davidson et al. 2017).

The genetic and developmental origins of variation in iris pigmentation are poorly understood. While iris color varies widely among species, variability in iris color is limited within most species (Negro et al. 2017). However, intraspecific variation in iris color is widespread in humans and certain domestic species (Negro et al. 2017). In mammals, iris colors typically include shades of brown, green, and blue. These colors all arise from varying concentrations and deposition patterns of melanin pigments in the iris (Edwards et al. 2016). In contrast, the diversity of eye colors in amphibians and birds also depends on the presence of non-melanin pigments. In birds, brilliant reds, oranges, and yellows arise from multiple non-melanin pigment types, including pteridines, purines, and carotenoids (Oliphant 1987a).

The domestic pigeon, *Columba livia*, shows intraspecific iris color variation among its 300+ different breeds. This variation, coupled with extensive genetic resources, makes the pigeon an ideal model to understand the genetics of iris pigmentation. Pigeons have three main iris colors: orange, pearl (white), and bull (dark brown; Fig. 1A). Orange iris color is the ancestral state (Bond 1919), and “orange” eyes in actuality range in shades from yellow to red, depending on the density of blood vessels in the eye (Hollander and Owen 1939; Sell 2012). The pearl iris color is white, with tinges of pink and red from blood vessels. Lastly, the bull iris color is named based on the similarity in color to dark bovine eyes, and ranges from dark brown to almost black (Hollander and Owen 1939; Levi 1986). Breeding experiments show that the switch between orange and pearl eye color is controlled by a single autosomal locus, and that orange is dominant to pearl (Bond 1919). Less is known about the inheritance of bull eye color. While orange and pearl irises can be found in a variety of pigeon breeds, the bull iris color is primarily found in birds with white plumage (Hollander, 1939). Breeders have also reported birds with a phenotype known as “odd eyes” (Levi 1986; Sell 2012), where one iris is a dark bull color and the other is either orange or pearl, suggesting that bull eye color may have a stochastic component.

**Figure 1.**
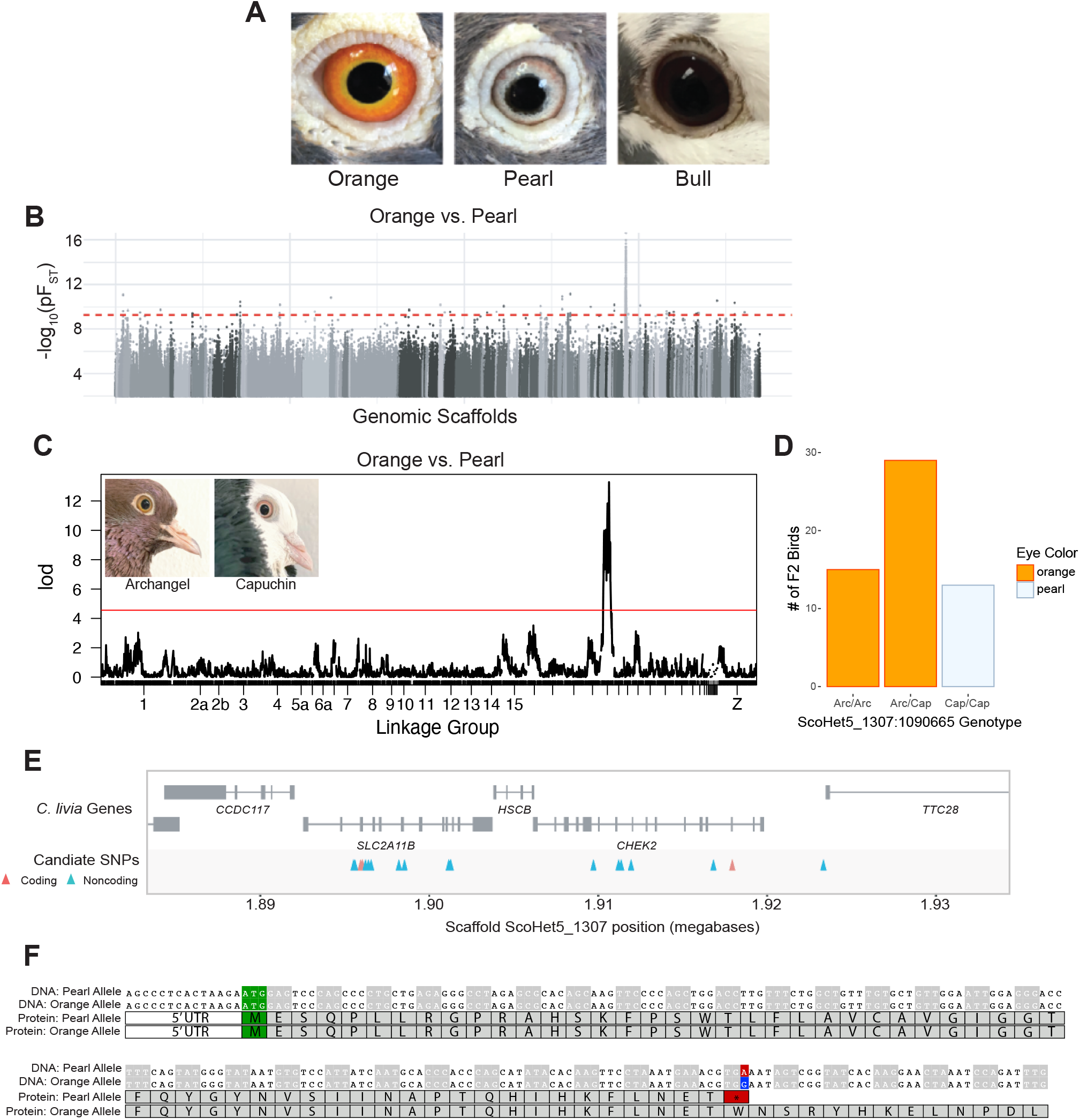
A single genomic locus is associated with pearl iris color in domestic pigeons. (A) Domestic pigeons typically have one of three major iris colors: the wild-type orange, pearl, or bull. (B) Whole-genome pF_ST_ comparisons of orange-eyed and pearl-eyed pigeons. Gray dots represent SNPs, with different shades indicating different genomic scaffolds. Dashed red line indicates genome-wide significance threshold. (C) Genome-wide QTL scan for pearl eye in the Archangel x Old Dutch Capuchin cross. Red line indicates 5% genome wide significance threshold. Insets: Archangel (left) and Capuchin (right) founders. (D) Eye color phenotypes of F_2_ progeny with different genotypes at the QTL peak marker. Arc, allele from the Archangel founder. Cap, allele from the Capuchin founder. (E) Genomic context of the pearl eye candidate region. Gene models for the region are shown in gray. SNPs in coding regions are shown in red, SNPs in non-coding regions are shown in blue. (F) Alignment of DNA (top) and predicted protein (bottom) sequences of SLC2A11B for pearl-eyed and orange-eyed pigeons. The start codon is highlighted in green. The DNA polymorphism at position ScoHet5_1307:1895934 is marked in red (pearl allele) or blue (orange allele); the resulting stop codon in the pearl allele is highlighted in red.

Pigeons have two types of non-melanin pigments in the iris, guanidines and pteridines (Oliphant 1987a). Guanidines are whitish opaque pigments, and pteridines are yellow-orange pigments (Oliphant 1987b). In orange-eyed pigeons, both guanidine and pteridine pigments are present in the iris stroma, while in white-eyed pigeons, only guanidine pigment is present (Oliphant 1987b; Oliphant 1987a). In bull-eyed pigeons, both white and orange pigments are absent from the iris stroma, so the underlying dark melanin pigment of the iris pigment epithelium is not obscured (Bond 1919; Oliphant 1987b). The genetic and developmental mechanisms underlying loss of pteridine iris pigment in pearl-eyed birds or both pteridine and guanidine pigment in bull-eyed birds are currently unknown. Loss could arise from defects in pigment production or failure to transport the pigment into the iris stroma, for example.

To better understand the genetic mechanisms that control iris color in domestic pigeons, we used a combination of genomic mapping and laboratory crosses to identify two loci that are associated with eye color. We identified a nonsense mutation that segregates with pearl eye color and a second locus associated with bull eye color. We also found a genetic link between iris and feather color in birds with an array of plumage depigmentation phenotypes collectively known as piebalding, thereby establishing a genetic link that explains the anecdotal co-occurance of iris and plumage color traits.

## RESULTS

### A single genomic locus is associated with pearl eye color in the domestic pigeon

To determine the genetic basis of pearl eye color, we compared whole-genome sequences from a diverse panel of orange-eyed (n = 28 from 17 domestic breeds and feral pigeons) and pearl-eyed pigeons (n = 33 from 25 breeds and ferals) using a probabilistic measure of allele frequency differentiation (pF_ST_; Domyan et al. 2016; Fig. 1B). We identified a single, 1.5-megabase genomic region on scaffold ScoHet5_1307 that was significantly differentiated between orange-eyed and pearl-eyed birds (ScoHet5_1307:1490703-3019601, genome wide significance threshold pF_ST_ = 5.4 × 10^−10^).

### The pearl eye locus segregates with eye color in F_2_ crosses

To confirm the association between the ScoHet5_1307 locus and pearl eye color, we turned to quantitative trait locus (QTL) mapping in an F_2_ intercross between an orange-eyed Archangel and a pearl-eyed Old Dutch Capuchin. Within this cross, F_2_ birds had either two pearl eyes (n=12), one pearl eye and one bull eye (n=1), two orange eyes (n=40), or one orange eye and one bull eye (n=5). We used a binary QTL model to compare birds with at least one pearl eye to birds with at least one orange eye, and identified a single peak on linkage group 20 associated with eye color (Fig. 1C, peak marker ScoHet5_1307: 1090556; peak LOD score = 13.28; candidate region, defined as a 2-LOD interval from the peak marker, spans ScoHet5_149:3706619 - ScoHet5_1307:1911647). The genotype at peak marker ScoHet5_1307:1090556 is perfectly associated with eye color in the cross, with all pearl-eyed F_2_ birds homozygous for the pearl-eyed Capuchin allele (Fig. 1D). We additionally used targeted genotyping in F_2_ individuals from a cross between Racing Homer and Parlor Roller breeds; this cross had four founders, one or more of which was heterozygous for the pearl allele so QTL mapping was impractical. We instead used PCR and Sanger sequencing to genotype a single nucleotide polymorphism (SNP) within the pearl-eye haplotype (ScoHet5_1307:1901234). This SNP again showed perfect association with the pearl eye phenotype (n = 25 F_2_ birds; *p* = 2.24 × 10^−7^, Fisher’s exact test). Thus, two independent approaches converged on the same genomic region controlling pearl eye.

### Pearl-eyed birds harbor a premature stop codon in solute carrier *SLC2A11B*

Because pearl eye color is recessive to orange, we searched for SNPs within the overlapping pF_ST_ and QTL peak region to identify polymorphisms where pearl-eyed birds were always homozygous for the reference allele (the Danish tumbler pigeon sequenced for the Cliv_2.1 reference assembly had pearl eyes; (Shapiro et al. 2013; Holt et al. 2018; Fig. S1A). We identified 20 SNPs spanning a 22-kb region (ScoHet5_1307:1895934-1917937) that showed the expected segregation pattern between orange-eyed and pearl-eyed birds, three of which were in protein coding regions, while 17 were intronic or intergenic (Fig. 1E). To evaluate variants in the pearl eye candidate locus, we first assessed the predicted impact of the three coding mutations identified in pearl-eyed birds. Two of these coding mutations are in *SLC2A11B*, a predicted solute carrier, while the third is in *CHEK2*, a kinase required for checkpoint-mediated cell-cycle arrest in response to DNA damage (Hirao et al. 2000; Chen et al. 2005). The coding mutation in exon 13 of *CHEK2* (position ScoHet5_1307:1917937) results in a synonymous substitution. *CHEK2* is not known to play a role in pigmentation or pteridine deposition; thus, this substitution is unlikely to have an effect on protein function or eye color phenotype.

The function of the second gene harboring coding mutations, *SLC2A11B*, is not well characterized, but is predicted to be a solute carrier. The first coding mutation in *SLC2A11B* (Position ScoHet5_1307: 1895934) changes a tryptophan (orange allele) to a premature stop codon (pearl allele; Fig. 1F). The second coding mutation (position ScoHet5_1307:1896042) results in a synonymous substitution 36 amino acids downstream of the premature stop codon. Unlike *CHEK2, SLC2A11B* is a strong candidate gene for pearl eye color. This gene has orthologs in fish and sauropsids, but not in mammals. Data from fish orthologs suggests that *SLC2A11B* plays a role in pigmentation. In medaka, for example, *SLC2A11B* is involved in the differentiation of pteridine-containing leucophore and xanthophore cells in scales (Kimura et al. 2014). The most closely related mammalian gene appears to be *SLC2A11* (*GLUT11*), a glucose transporter (Doege et al. 2001; Kimura et al. 2014).

The premature stop codon in pearl-eyed pigeons falls in exon 3 of *SLC2A11B*, which is predicted to severely truncate the resulting protein from 504 to 57 amino acids. Translation initiation at the next in-frame methionine would produce a protein missing the first 95 amino acids, but with the remaining 81% (409 out of 504 amino acids) of the protein intact. To predict if such a truncated protein would be functional, we used InterProScan (Zdobnov and Apweiler 2001) and Phobius (Käll et al. 2004) to predict transmembrane domains and conserved functional motifs within the SLC2A11B protein, and examined sequence similarity across species (Fig. S1 B-D). The first 94 amino acids of SLC2A11B are predicted to code for two transmembrane domains that are highly conserved; removing these domains is predicted to be detrimental to protein function (PROVEAN score of -189.145; Choi et al. 2012; Choi and Chan 2015). Therefore, the pearl mutation, which results in a loss of pteridines in the iris, is predicted to truncate a highly-conserved protein that is associated with the differentiation of pteridine-containing pigment cells. The first two transmembrane domains of the SLC2A11B protein are highly conserved across species, yet we identified orthologs in two bird species, hooded crow [NCBI accession XP_019140832.1] and wire-tailed manakin [NCBI accession XP_027569903.1], in which the annotated protein sequence is missing the first of these transmembrane domains. While we cannot rule out a misannotation in these genomes, neither species appears to have yellow-orange iris pigment. Hooded crows have dark eyes, while wire-tailed manakins have white irises (Madge 2020; Snow 2020), raising the possibility that neither species is capable of producing pteridine iris pigments due to a hypomorphic or null version of SLC2A11B.

### Expression of the *SLC2A11B* pearl allele is reduced

Using high-throughput RNA-sequencing (RNA-seq) datasets, we found that *SLC2A11B* shows very low levels of expression in most adult tissues, including the retina, but substantial expression in both Hamburger-Hamilton stage 25 (HH25; Hamburger and Hamilton 1951) whole embryos (n = 2) and embryo heads (n = 12) (Fig. S2 A-B). Based on genotypes at the two coding SNPs in the pearl eye haplotype (nonsense SNP ScoHet5_1307:1895934 and synonymous SNP ScoHet5_1307:1896042), we found that embryo head samples homozygous for the pearl allele show a significant reduction in *SLC2A11B* expression (*p* = 3.94 × 10^−6^, two-tailed t-test; Fig. S2C). Analysis of read distribution within the *SLC2A11B* gene shows a decrease in spliced reads specifically within the first three annotated exons, suggesting that alternative splicing or nonsense-mediated decay may be occurring. In summary, the pearl eye phenotype is associated with a nonsense mutation in a known mediator of yellow-orange pigments, which in turn is linked to a significant decrease in *SLC2A11B* expression, possibly due to nonsense-mediated decay of the mutant transcripts.

### QTL mapping identifies a single genomic locus associated with bull eye color

Variation at *SLC2A11B* appears to act as a switch between two of the major pigeon iris colors, orange and pearl eye, but it does not explain the third major color, bull eye. Bull eyes are dark brown, lacking both orange and white pigment in the iris. However, pigeon breeders observe that bull eye color can occur on either an orange or pearl genetic background (Sell 2012), suggesting that the loss of orange pigment in bull eyes likely arises from a mechanism that does not involve *SLC2A11B*.

To identify the genetic basis of bull eye color, we used QTL mapping in two independent F_2_ intercrosses. In a cross between an orange-eyed Pomeranian Pouter and a bull-eyed Scandaroon, F_2_ birds had either two bull eyes (n=41), two orange eyes (n=40), or “odd eyes”, where one eye has a pigmented iris stroma and the other eye is bull (n=12) (Fig. 2A). Using a binary model where odd-eyed birds were included in the “bull eye” group, we identified a QTL on linkage group 15 (Fig. 2C; peak marker ScoHet5_507: 11175287, LOD score = 11.89, genome wide significance threshold = 4.28). The peak region spans 2.0 Mb across two genomic scaffolds, from ScoHet5_507:9736663 to scaffold ScoHet5_683.1: 279252, and includes 42 annotated genes. Nearly all (52/53) odd-eyed or bull-eyed F_2_ birds have at least one copy of the bull-eyed Scandaroon allele at the peak marker (Fig. 2D), indicating dominant inheritance of bull eye color. However, penetrance is both incomplete and lower in heterozygotes, suggesting a stochastic effect of the bull eye allele. The one odd-eyed bird that is homozygous for the orange Pomeranian Pouter allele may be a recombinant between the peak QTL marker and the causative bull eye variant.

**Figure 2.**
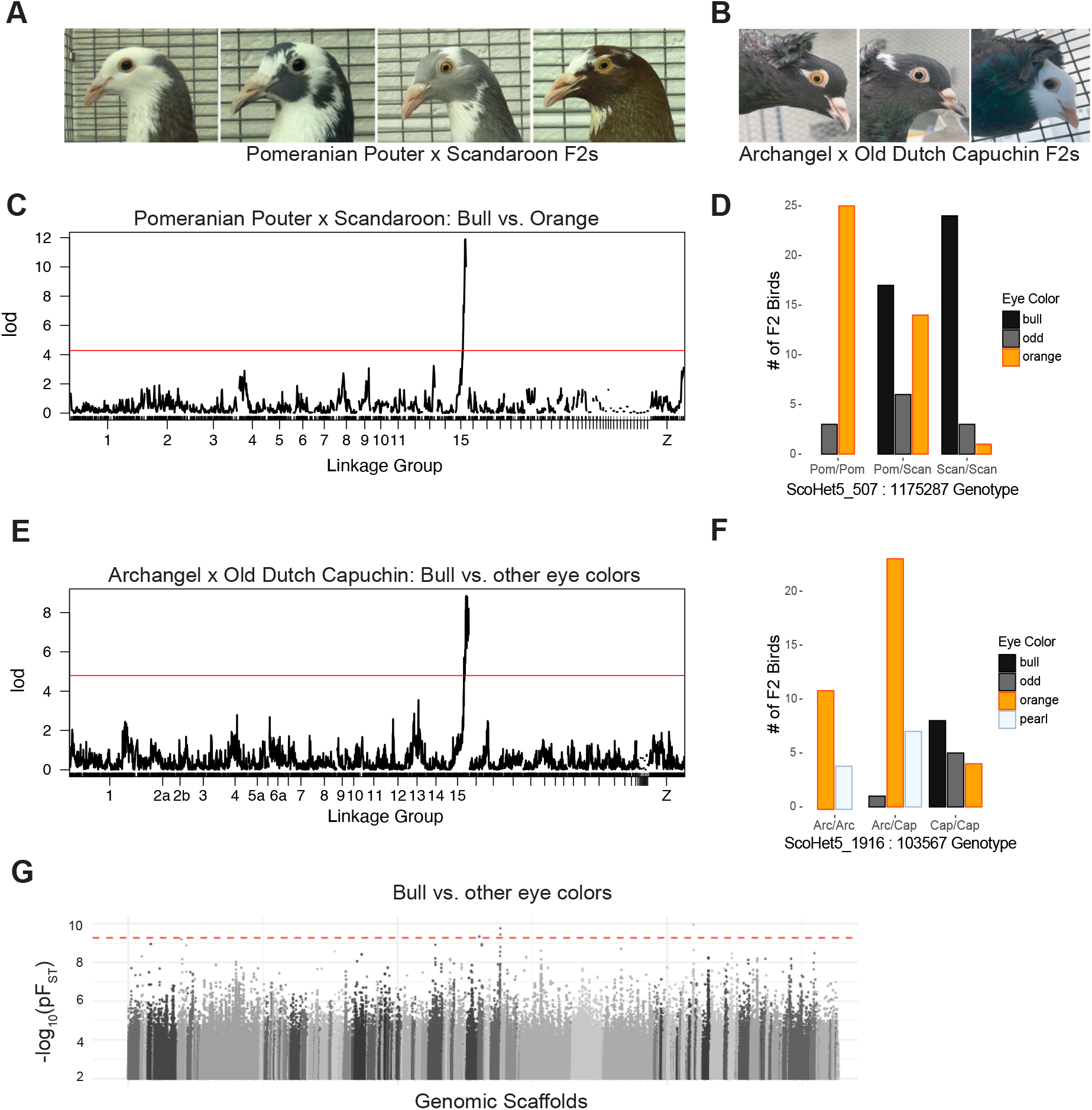
A single genomic locus is associated with bull eye color in two F_2_ intercrosses. **(A)** F_2_ offspring from an intercross between a Pomeranian Pouter and a Scandaroon have either bull (left two images) or orange (right two images) eyes. **(B)** F_2_ offspring from an intercross between an Archangel and an Old Dutch Capuchin have orange (left), pearl (center), or bull (right) eyes. **(C)** Genome-wide QTL scan of the Pomeranian Pouter x Scandaroon cross for bull eye. Red line indicates 5% genome wide significance threshold. **(D)** Iris color phenotype counts for each genotype at the bull eye peak marker from the Pomeranian Pouter x Scandaroon cross. Pom, allele from Pomeranian Pouter founder. Scan, allele from Scandaroon founder. **(E)** Genome-wide QTL scan of the Archangel x Old Dutch Capuchin cross for bull eye. Red line indicates 5% genome wide significance threshold. **(F)** Iris color phenotype counts for each genotype at the bull eye peak marker from the Archangel x Capuchin cross. Arc, allele from the Archangel founder. Cap, allele from the Capuchin founder. **(G)** Whole-genome pF_ST_ comparisons of bull-eyed birds to birds with non-bull (orange or pearl) eyes. Dashed red line indicates 5% threshold for genome-wide significance.

In the cross between the orange-eyed Archangel and the pearl-eyed Old Dutch Capuchin that we originally used to map pearl eyes, neither founder had the bull eye phenotype. However, some offspring had either two bull eyes (n = 8) or odd eyes (n = 6) (Fig. 2B). We used a binary model to compare these 14 birds with at least one bull eye to 52 F_2_ birds without bull eyes (either two orange or two pearl eyes). Here, too, we identified a single locus associated with bull eye color on linkage group 15 (Fig. 2E; peak marker ScoHet5_1916:103567, peak LOD score = 8.85, genome wide significance threshold = 4.61). The peak region spans 1.5 Mb across eight scaffolds, including the same two scaffolds identified in the Scandaroon x Pomeranian Pouter cross, and captures 44 annotated genes. Although the Old Dutch Capuchin founder does not have bull eyes, nearly all bull-eyed and odd-eyed F_2_s carry two copies of the Capuchin allele at the peak marker (Fig. 2F). This suggests that, unlike in the Pomeranian Pouter x Scandaroon cross, inheritance of bull eye color in the Archangel x Capuchin cross is recessive with low penetrance. The lone odd-eyed bird in the latter cross is heterozygous for the Capuchin allele at ScoHet5_1916:103567 and may have a recombination event between the peak QTL marker and the causative bull eye variant.

### Bull eye color and allelic heterogeneity

QTL mapping identified a single locus associated with bull eye color in two different crosses, but the inheritance pattern of bull eye appears to differ in each case. Furthermore, genome-wide pF_ST_ analysis comparing bull-eyed birds (n = 18) to a background dataset of both orange-eyed and pearl-eyed birds (n = 61) identified a small number of differentiated SNPs across multiple scaffolds, including ScoHet5_507, but did not pinpoint a single well-differentiated region in all bull-eyed breeds (Fig. 2G). Together, these results imply that, while our QTL mapping identified the same genomic region in two separate crosses, the variants that give rise to bull eye color are probably not the same across all pigeon breeds.

### Bull eye color is associated with plumage depigmentation

Pigeon hobbyists have long noted that bull eye color is most common in birds with white plumage (Sell 2012), including individuals with solid white plumage and those with a range of piebalding phenotypes. Piebalding is characterized by broad regions of white and pigmented feathers, and these regionalized de-pigmentation patterns are often breed-specific. Both the Scandaroon and Pomeranian Pouter cross founders show breed-specific piebald patterning, and the F_2_ offspring of this cross show highly variable piebalding across multiple body regions (Fig. 3A-B). We found that plumage color in many body regions is significantly associated with bull eye color in the Pomeranian Pouter x Scandaroon cross. The strength of this relationship varies by region, with areas like the lateral head and dorsal wing (Fig. 3C-D) showing a stronger relationship with eye color than the lateral neck (Fig. 3E; additional body regions are shown in Fig. S3).

**Figure 3.**
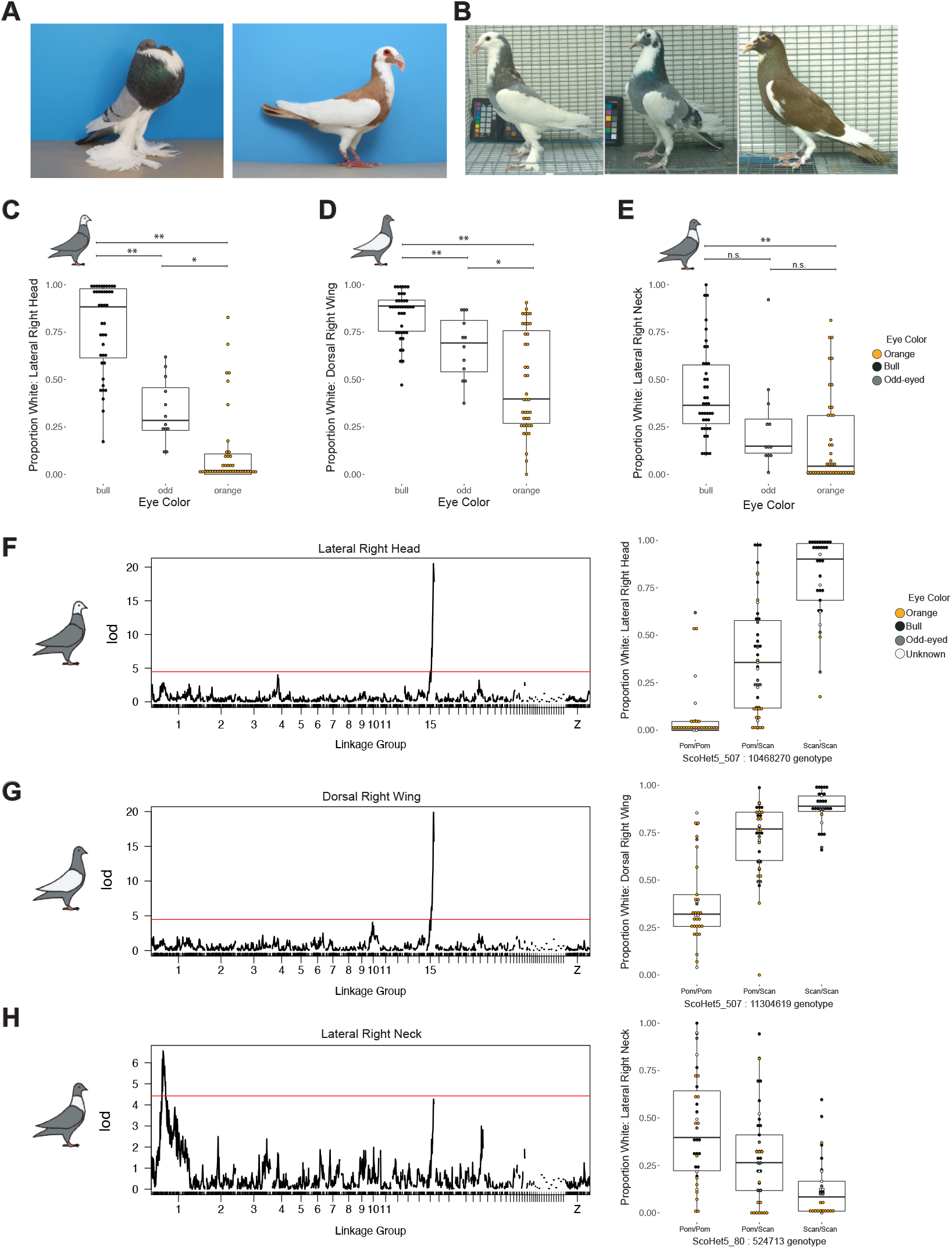
Bull eye color is associated with white plumage in an F_2_ intercross. (A) Examples of standard plumage patterning for the Pomeranian Pouter (left) and Scandaroon (right) breeds. Photos by Layne Gardner, used with permission. (B) Examples of variable piebald plumage patterning in Pomeranian Pouter x Scandaroon F_2_ offspring. (C-E) Boxplots of association between eye color and proportion of white plumage on the (C) lateral right head, (D) dorsal right wing, and (E) lateral right neck of F_2_ birds. **, *p* ≤0.0001; *, 0.001 < *p* ≤ 0.01; n.s., *p* >0.01. Boxes span from the first to third quartile of each data set, with lines indicating the median. Whiskers span up to 1.5x the interquartile range. (F-H) QTL scans for proportion of white plumage (left side of panel) and proportion of white plumage by genotype at the peak marker (right) for (F) lateral right head, (G) dorsal right wing, and (H) lateral right neck. Red line, 5% genome-wide significance threshold.

To further evaluate the genetic relationship between piebalding and bull eye color, we quantified the proportion of white plumage across 15 different body regions in the F_2_ progeny of the Pomeranian Pouter x Scandaroon cross. We identified two broad QTL regions associated with white plumage (Fig. 3F-H, Fig. S4). Each locus is associated with white plumage in specific body regions and explains 15-58% of the variance in the cross (Table 1). The QTL on linkage group 1 is associated with white plumage on the neck and dorsal body, and individuals with white plumage carry the Pomeranian Pouter allele at the linkage group 1 candidate locus. The QTL on linkage group 15 is associated with white plumage on the head, wings, and dorsal body; for this locus, the Scandaroon allele is associated with white plumage. The linkage group 15 piebalding QTL overlaps with the locus identified for bull eye, suggesting either closely linked variants in the same or different genes, or the same gene controlling both traits. These associations are consistent with breed-specific plumage patterns, as Scandaroon pigeons typically have white plumage on the head, wings, and ventral body, while the Pomeranian Pouter breed is characterized by a white “bib” on the neck (see examples, Fig. 3A). In summary, at least two genetic loci control piebalding in pigeons, one of which overlaps with the bull eye locus, and these loci act in a regionally- and breed-specific manner.

**Table 1.** Summary of QTL for regional white plumage. PVE, percent variance explained; Pom, Pomeranian Pouter; Scan, Scandaroon.

| Body Region | Linkage Group | Peak Marker | Peak LOD score | PVE | Associated Allele |
| --- | --- | --- | --- | --- | --- |
| Dorsal Right Wing | 15 | ScoHet5_507_11304619 | 19.9 | 57 | Scan |
| Dorsal Left Wing | 15 | ScoHet5_507_11304619 | 15.5 | 49 | Scan |
| Dorsal Body | 1 | ScoHet5_80_11511249 | 9.11 | 30 | Pom |
| Dorsal Body | 15 | ScoHet5_683.1_42424 | 4.45 | 15 | Scan |
| Dorsal Neck | 1 | ScoHet5_80_3402497 | 5.72 | 22 | Pom |
| Dorsal Head | 15 | ScoHet5_507_11175287 | 22.4 | 63 | Scan |
| Ventral Right Wing | 15 | ScoHet5_507_11175287 | 12.2 | 40 | Scan |
| Ventral Left Wing | 15 | ScoHet5_507_11227444 | 9.34 | 33 | Scan |
| Ventral Body | 15 | ScoHet5_507_11058018 | 11.4 | 38 | Scan |
| Ventral Tail | 15 | ScoHet5_683.1_42424 | 17.4 | 52 | Scan |
| Ventral Neck | 1 | ScoHet5_80_524713 | 4.81 | 21 | Pom |
| Ventral Neck | 15 | ScoHet5_507_11175287 | 7.39 | 30 | Scan |
| Lateral Right Head | 15 | ScoHet5_507_10468270 | 20.5 | 58 | Scan |
| Lateral Right Neck | 1 | ScoHet5_80_524713 | 6.56 | 15 | Pom |
| Lateral Left Head | 15 | ScoHet5_507_11058018 | 20.7 | 58 | Scan |
| Lateral Left Neck | 1 | ScoHet5_2444_504541 | 4.37 | 17 | Pom |

### *EDNRB2* is a candidate gene for bull eye color and white plumage

We next wanted to identify candidate genes for bull eye color and white plumage within the linkage group 15 region. Of the 60 genes included in at least one of the Pomeranian Pouter x Scandaroon (2.0 Mb, 42 genes) or Archangel x Old Dutch Capuchin (1.5 Mb, 44 genes) bull eye QTL peaks, comparison to gene ontology databases did not identify any genes with GO annotations related to pigmentation. However, we were able to find potential links to pigment patterning for five genes, including the endothelin receptor *EDNRB2* (Table 2). Mutations in *EDNRB2* are associated with depigmentation phenotypes in several domestic bird species, including “Panda” plumage in Japanese Quail, spot patterning in ducks, tyrosinase-independent mottling in chickens, and white plumage with dark eye color in Minohiki chickens (Miwa et al. 2007; Kinoshita et al. 2014; Li et al. 2015; Xi et al. 2020). Additionally, changes in the mammalian orthologue *ENDRB* are responsible for piebalding phenotypes in mice and the piebald-like frame overo pattern in horses (Koide et al. 1998; Metallinos et al. 1998). Given the known role of endothelin receptors in piebalding in other vertebrates, *EDNRB2* is a compelling candidate for the linked piebalding and bull eye phenotypes in domestic pigeons. We examined the allele frequencies and genotypes of SNPs within *EDNRB2* coding regions in both the bull-eyed and non-bull-eyed populations used for pF_ST_ analysis and did not identify any coding polymorphisms that were unique to bull-eyed birds, suggesting that noncoding regulatory changes may mediate bull eye color and piebalding in domestic pigeons. Due to the allelic heterogeneity and incomplete penetrance of the bull eye phenotype, however, we cannot rule out coding changes in *EDNRB2*, or other candidate genes within the region, as mediators of the bull eye phenotype.

**Table 2.** Summary of pigment-associated genes within the LG15 QTL.

| Gene Name | Scaffold | Role in Pigmentation |
| --- | --- | --- |
| *CDX1* | ScoHet5_507 | Involved in neural crest development, Reduction in *CDX1* is associated with white spotting in mice (Sanchez-Ferras et al. 2016) |
| *NSDHL* | ScoHet5_507 | Mice with heterozygous mutations can have striped coats (Liu et al. 1999) |
| *VAMP7* | ScoHet5_507 | SNARE protein involved in TYRP1 trafficking to the melanosome (Tamura et al. 2011) |
| *EDNRB2* | ScoHet5_507 | Controls migration of neural crest derived pigment cells (Pla et al. 2005; Harris et al. 2008); Linked to plumage pigmentation phenotypes in multiple avian species (Miwa et al. 2007; Kinoshita et al. 2014; Li et al. 2015; Wu et al. 2017; Xi et al. 2020) |
| *GPR119* | ScoHet5_1916 | G-protein coupled receptor expressed in human melanocytes (Scott et al. 2006) |

## DISCUSSION

### *SLC2A11B* and pearl eyes

Using comparative genomic and classical genetic approaches, we identified two candidate loci that control the three major iris colors of domestic pigeons. A locus on scaffold ScoHet5_1307 is associated with pearl eye color. This region contains a SNP fixed in pearl-eyed birds that changes a tryptophan to a premature stop codon in exon 3 of the solute carrier *SLC2A11B*, and was also recently identified by Andrade *et al*. and Si *et al*. as a candidate mutation for pearl eye color in pigeons (Si et al. 2020; Andrade et al. 2021). We found that the nonsense mutation is associated with pearl iris color in individually phenotyped pigeons from a wide array of domestic breeds, consistent with a single mutation arising early in domestication (Si et al. 2020). We also showed that the *SLC2A11B* locus is the one and only genomic region that segregates with pearl eye color in two F_2_ crosses. Our results support the trio genotyping of the *SLC2A11B* mutation performed by Andrade *et al*. (Andrade et al. 2021), and our linkage mapping excludes a role for the remainder of the genome in the switch between orange and pearl eyes. Intriguingly, while all pearl-eyed birds in our sample share a common *SLC2A11B* allele, pigeon breeders have also identified a second locus associated with white iris color that appears to be genetically distinct and is linked to brown plumage color (Levi 1986; Sell 2012). Future analysis of individual birds with this “false pearl” eye color could expand our understanding of the genes affecting pteridine synthesis and localization in the eyes of birds.

The *SLC2A11B* gene is not well-characterized, but likely plays an evolutionarily conserved role in the development of pteridine-containing pigment cells. A nonsense mutation in *SLC2A11B* in medaka is associated with loss of mature pteridine-containing leucophores and xanthophores, and the Zebrafish Mutation Project identified differentiation defects in *SLC2A11B*-mutant xanthophores (Kimura et al. 2014). Si *et al*. (2020) additionally identified a frameshift mutation in *SLC2A11B* in cormorants, which have unique blue eyes and appear to lack pteridine pigments in the iris. Similarly, the missing transmembrane domain in the manakin and crow described here might render SLC2A11B incapable of pteridine deposition.

*SLC2A11B* does not have a mammalian ortholog, and its presence is restricted to species that have xanthophores or xanthophore-like cells (Kimura et al. 2014). Comparative analysis of solute carriers across species shows that the *SLC2A11B* gene likely originated prior to the teleost fish-specific genome duplication, and was then lost in mammals (Kimura et al. 2014). Loss of *SLC2A11B* may have restricted the repertoire of pigments that mammals can synthesize.

### Allelic heterogeneity at the bull eye locus

Observations by pigeon breeders previously indicated a simple recessive mode of inheritance for pearl eye color (Sell 2012), and this is confirmed by our analyses. The third major iris color in domestic pigeons, bull eye, appears to have a more complicated inheritance pattern. Through QTL mapping in two F_2_ crosses, we identified a single genomic locus on linkage group 15 that is associated with bull eye color. As previously noted by breeders, bull eye color is associated with white plumage (Sell 2012), and QTL mapping identified a strong association between the same linkage group 15 locus and piebald plumage patterning on the wing and head.

Despite the overlap in QTL for bull eye color in two F_2_ crosses and the QTL white plumage, we were unable to pinpoint a single mutation within this locus associated with bull eye color through a comparative genomic approach. This suggests that bull eye may not be caused by a single genetic variant that is shared across breeds. Instead, the linkage group 15 QTL regions may harbor multiple breed-specific mutations. These mutations may affect multiple closely linked genes or may impact a single gene. Future work will examine the genetic underpinnings of regionalized plumage patterning in F_2_ crosses and work towards identification of specific genetic variants associated with bull eye color and the piebald plumage that typically accompanies it.

### *EDNRB2* and constraints on endothelin receptor evolution

Although the specific mutations that cause bull eye color and white plumage color remain unknown, the linkage group 15 QTL for bull eye color and piebalding contains a strong candidate gene, *EDNRB2*. The endothelin signaling pathway plays critical roles in the development and migration of multiple neural crest cell populations, including pigment cells. In mammals, mutations in the endothelin receptor *ENDRB* are linked to piebalding in mice; lethal white foal syndrome in horses; and Waardenburg Shah syndrome type 4A in humans, which is characterized by changes in hair, skin, and eye pigment, as well as congenital defects in enteric nervous system development (Read and Newton 1997; Koide et al. 1998; Metallinos et al. 1998; Jabeen et al. 2012). In several bird species, coding and regulatory variants of *EDNRB2* are associated with white plumage phenotypes and dark eye color (Miwa et al. 2007; Kinoshita et al. 2014; Li et al. 2015; Wu et al. 2017; Xi et al. 2020), but they are not typically linked to other major pathologies. Thus, while endothelin signaling is linked to pigmentation changes across vertebrates, *ENDRB* mutations in mammals are typically associated with deleterious pleiotropic effects, while *EDNRB2* mutations in birds are not.

The endothelin signaling pathway in vertebrates evolved through multiple rounds of gene duplication, and most bony vertebrates have three endothelin receptor genes: *EDNRA, EDNRB1*, and *EDNRB2* (Braasch, Volff, et al. 2009). Expression of different combinations of endothelin receptors and ligands characterize unique cell populations. In *Xenopus*, chicken, and quail, for example, *EDNRB2* is expressed specifically in migrating and post-migratory melanophores, while non-pigment neural crest populations, like skeletal and trunk neural crest cells, express *EDNRA* or *EDNRB1* (Square et al. 2016). However, *EDNRB2* was lost in therian mammals, and the sole endothelin B receptor *ENDRB* is expressed in both trunk neural crest populations and melanophores (Braasch, Volff, et al. 2009; Square et al. 2016). As a result, in therian mammals, changes in endothelin signaling typically affect both pigmentation and neurogenesis. The retention of *EDNRB2* in non-mammalian vertebrates, on the other hand, may permit the evolution and development of novel pigment patterns because the genetic controls of pigment cell migration and neurogenesis are uncoupled.

### Gene duplication and retention mediate the evolution of pigment diversity

The retention of *EDNRB2* in non-mammalian vertebrates, and the diverse endothelin-mediated pigment patterns identified across bird species, point to a role for gene duplication in mediating or constraining diversity in both pigment type and patterning. In species that retained *EDNRB2*, sub-functionalization mediates the evolution of novel pigment patterns such as piebalding, while in species that lost *EDNRB2*, such changes are severely constrained by the requirement for a functional endothelin receptor B gene. This idea of gene loss restricting pigment phenotypes is also relevant to the retention of our pearl eye candidate gene *SLC2A11B*, which is only present in species with pteridine-containing xanthophore- or leucophore-like cells. Solute carriers in the *SLC2A* family also evolved through multiple rounds of gene duplication, though their evolutionary history is not as clear as that of endothelin ligands and receptors due to multiple segmental duplication events (Kimura et al. 2014; Lorin et al.2018). Gene duplication and retention permitted the striking expansion and evolution of novel pigment types and patterns in teleost fish (Braasch, Brunet, et al. 2009; Lorin et al. 2018). The identification of *SLC2A11B* and *EDNRB2* as candidate genes for pigeon eye color suggests that similar patterns of retention of gene duplicates may mediate the evolution of pigment phenotypes across vertebrate species.

## MATERIALS AND METHODS

### Animal husbandry and phenotyping of F_2_ offspring

Pigeons were housed in accordance with protocols approved by the University of Utah Institutional Animal Care and Use Committee (protocols 10-05007, 13-04012, and 19-02011). Two intercrosses were used in these studies. An intercross between a male Pomeranian Pouter and two female Scandaroons was performed to generate 131 F_2_ offspring (Domyan et al. 2016). An intercross between a male Archangel and a female Old Dutch Capuchin generated 98 F_2_ offspring.

### Whole Genome Resequencing

DNA was extracted from blood samples collected with breeders’ written permission at the annual Utah Premier Pigeon Show or from our lab colony using the Qiagen DNEasy Blood and Tissue Kit (Qiagen, Valencia, CA). Samples were treated with RNAse during extraction. Isolated DNA was submitted to the University of Utah High Throughput Genomics Shared Resource for library preparation using the Illumina Tru-Seq PCR-Free library kit. The resulting libraries were sequenced on either the Illumina HiSeq or Illumina NovaSeq platforms. Raw sequence data for 54 previously unpublished samples is available in the NCBI Sequence Read Archive under BioProject accession PRJNA680754. These data sets were combined with previously published data sets (BioProject accessions PRJNA513877, PRJNA428271, and PRJNA167554) for variant calling.

### Genomic Analyses

Variant calling was performed with FastQForward, which wraps the BWA short read aligner and Sentieon (<u>sentieon.com</u>) variant calling tools to generate aligned BAM files (fastq2bam) and variant calls in VCF format (bam2gvcf). Sentieon is a commercialized variant calling pipeline that allows users to follow GATK best practices using the Sentieon version of each tool (<u>broadinstitute.org/gatk/guide/best-practices</u> and <u>support.sentieon.com/manual/DNAseq_usage/dnaseq/</u>). FastQForward manages distribution of the workload to these tools on a compute cluster to allow for faster data-processing than when calling these tools directly, resulting in runtimes as low as a few minutes per sample.

Raw sequencing reads from the 54 resequenced individuals described above were aligned to the Cliv_2.1 reference assembly (Holt et al. 2018) using fastq2bam. Variant calling was performed for each newly resequenced individual, as well as 132 previously resequenced individuals (Shapiro et al. 2013; Domyan et al. 2016; Vickrey et al. 2018; Bruders et al. 2020), using bam2gvcf and individual genome variant call format (gVCF) files were created. Joint variant calling was performed on a total of 186 individuals using the Sentieon GVCFtyper algorithm. The resulting VCF file was used for all subsequent genomic analyses.

The subsequent variant call format (VCF) file was used for pF_ST_ analysis using the GPAT++ toolkit within the VCFLIB software library (https://github.com/vcflib). For orange vs. pearl pF_ST_ analysis, the genomes of 28 orange-eyed birds from were compared to the genomes of 33 pearl-eyed birds. For bull eye vs. other color pF_ST_ analysis, the genomes of 18 bull eyed birds were compared to the genomes of 61 non-bull birds (a mix of orange and pearl)

### Eye color phenotyping

Eye colors of birds in our whole genome resequencing panel were determined from photographs taken at the time of sampling. Each photograph was independently scored by three individuals. In instances where eye color could not confidently be determined from photographs, those individuals were not included in pF_ST_ analysis. Breeds included in the orange-eyed group: American Show Racer, Archangel, Chinese Owl, Damascene, Dragoon, English Carrier, Feral, Granadino Pouter, Hamburg Sticken, Hungarian Giant House Pigeon, Italian Owl, Mindian Fantail, Modena, Pomeranian Pouter, Rafeno Pouter, Saxon Pouter, and Starling. Breeds included in the pearl-eyed group: Australian Tumbler, Bacska Tumbler, Berlin Long Faced Tumbler, Berlin Short Faced Tumbler, Birmingham Roller, Budapest Tumbler, Chinese Owl, Cumulet, Danzig Highflier, English Short Faced Tumbler, English Trumpeter, Feral, Helmet, Indian Fantail, Long Face Clean Leg Tumbler, Naked Neck, Oriental Roller, Polish Lynx, Russian Tumbler, Saint, Temeschburger Schecken, Turkish Tumbler, Uzbek Tumbler, and Vienna Medium Faced Tumbler. Breeds included in the bull-eyed group: African Owl, Canario Cropper, Classic Old Frill, Chinese Nasal Tuft, Fairy Swallow, Ice Pigeon, Komorner Tumbler, Lahore, Mookee, Old German Owl, Oriental Frill, Scandaroon, and Schalkaldener Mohrenkopf. Eye colors of 93 Pomeranian Pouter x Scandaroon and 66 Archangel x Capuchin F_2_ birds were recorded based on observation at the time of euthanasia, and live photographs showing eye color were taken for reference.

### Plumage phenotyping

Following euthanasia, photos were taken of F_2_ plumage including dorsal and ventral views with wings and tail spread, and lateral views with wings folded. We divided the body into 15 different regions for phenotyping: dorsal head, right lateral head, left lateral head, dorsal neck, ventral neck, right lateral neck, left lateral neck, dorsal body, ventral body, dorsal tail, ventral tail, dorsal right wing, dorsal left wing, ventral right wing, and ventral left wing. To score each region, we imported photos into Photoshop v21.1.0×64 (Adobe, San Jose, CA) and used the magic wand tool to select only the white feathers within the body region. Following this selection, we saved two separate images: one containing the entire region (both pigmented and white feathers) with the color for the white feathers inverted (hereafter, “whole region image”), and one with the selected white feathers removed and only the pigmented feathers remaining (“pigmented region image”). For each body region, we imported these two images into ImageJ (v1.52a; Schneider et al. 2012) and converted them to greyscale, then used the threshold tool to measure the number of pixels in each image. To calculate the proportion of white feathers for each region, we subtracted the number of pixels in the pigmented region image from the number of pixels in the whole region image, then divided by the number of pixels in the whole region image.

### Genotype by Sequencing

DNA samples from founders of the crosses and their F_2_ progeny were extracted using the Qiagen DNeasy Blood and Tissue kit. Our Genotype by Sequencing approach was adapted from a previously published protocol with minor modifications (Elshire et al. 2011; Domyan et al. 2016). DNA was digested with ApeKI, and size selected for fragments in the 550-650 bp range. Domyan et al. (2016) performed an initial round of genotyping for the Pomeranian Pouter x Scandaroon cross. These libraries were sequenced using 100- or 125 bp paired-end sequencing on the Illumina HiSeq2000 platform at the University of Utah Genomics Core Facility. Genotype by sequencing for the Archangel x Capuchin founders (n=2) and F_2_ offspring (n=98), as well as supplemental sequencing for 20 additional and 17 previously low-coverage Pomeranian Pouter x Scandaroon F_2_s, was performed by the University of Minnesota Genomics Center. New GBS libraries were sequenced on a NovaSeq 1×100 SP FlowCell.

### Linkage Map Construction

Genotype by Sequencing reads were trimmed using CutAdapt (Martin 2011), then mapped to the Cliv_2.1 reference genome reads using Bowtie2 (Langmead and Salzberg 2012). Genotypes were called using Stacks2 by running “refmap.pl” (Catchen et al. 2013). In the Pomeranian Pouter x Scandaroon cross, which had three founders, the Pomeranian Pouter and one of the two Scandaroons designated as parents; to account for the three-founder cross structure, all markers where the genotypes of the two Scandaroon founders differed were subsequently removed from the dataset.

We constructed genetic maps using R/qtl v1.46-2 (www.rqtl.org; Broman et al. 2003). Autosomal markers showing significant segregation distortion (*p* < 0.01 divided by the total number of markers genotyped, to correct for multiple testing) were eliminated. Sex-linked scaffolds were assembled and ordered separately, due to differences in segregation pattern for the Z chromosome. Z-linked scaffolds were identified by assessing sequence similarity and gene content between pigeon scaffolds and the Z chromosome of the annotated chicken genome (Ensembl Gallus_gallus-5.0).

Pairwise recombination frequencies were calculated for all autosomal and Z-linked markers. Markers with identical genotyping information were identified using the “findDupMarkers” command, and all but one marker in each set of duplicates was removed. Within individual Cliv_2.1 scaffolds, markers were filtered by genotyping rate; to retain the maximal number of scaffolds in the final map, an initial round of filtering was performed to remove markers where fewer than 50% of birds were genotyped. Large scaffolds (> 40 markers) were subsequently filtered a second time to remove markers where fewer than 66% of birds were genotyped.

Within individual scaffolds, R/Qtl functions “droponemarker” and “calc.errorlod” were used to assess genotyping error. Markers were removed if dropping the marker led to an increased LOD score, or if removing a non-terminal marker led to a decrease in length of >10 cM that was not supported by physical distance. Individual genotypes were removed if they had error LOD scores >5 (a measure of the probability of genotyping error, see (Lincoln and Lander 1992). Linkage groups were assembled from both autosomal markers and Z-linked markers using the parameters (max.rf 0.15, min.lod 6). Scaffolds in the same linkage group were manually ordered based on calculated recombination fractions and LOD scores. Linkage groups in the Pomeranian Pouter x Scandaroon map were numbered by marker number. Linkage groups in the Archangel x Old Dutch Capuchin map were numbered based on scaffold content to correspond with Pomeranian Pouter x Scandaroon linkage groups.

### Quantitative Trait Locus Mapping

We performed QTL mapping using R/qtl v1.46-2 (Broman et al. 2003). For eye color phenotypes, we used the *scanone* function to perform a single-QTL genome scan using a binary model. In QTL scans for the bull eye phenotype, “odd-eyed” birds with a single bull eye were scored as bull. For piebalding phenotypes, we used the *scanone* function to perform a single-QTL genome scan using Haley-Knott regression. For each phenotype, the 5% genome-wide significance threshold was calculated by running the same *scanone* with 1000 permutation replicates. For each significant QTL peak, we calculated 2-LOD support intervals using the *lodint* function. We calculated percent variance explained (PVE) using the *fitqtl* function.

### *SLC2A11B* mutation identification and gene re-annotation

We identified numerous SNPs with maximal pF_ST_ scores, and manually examined genotype calls from the VCF file to identify SNPs that followed the expected recessive inheritance pattern of pearl eye (i.e., all pearl-eyed birds were homozygous for the reference allele and all orange-eyed birds were either heterozygous or homozygous for the alternate allele). We identified SLC2A11B orthologs across species using NCBI blastp (https://blast.ncbi.nlm.nih.gov/Blast.cgi; (Altschul et al. 1990; Johnson et al. 2008). The first 10-20 amino acids of the SLC2A11B protein vary across species, but alignments showed that the annotated pigeon protein was missing >80 amino acids that are well conserved most other species, and was likely incomplete. We then took RNA sequences for the orange and pearl alleles of *SLC2A11B* and translated each using Expasy Translate (https://web.expasy.org/translate/; (Gasteiger et al. 2003). The longest contiguous protein predicted for the pearl allele matched the protein sequence available on NCBI, while the longest contiguous protein for the orange allele was in the same open reading frame, but contained an additional 95 amino acids at the start of the protein sequence. This N-terminal sequence matched the highly conserved SLC2A11B protein sequence annotations across species. The amino acid residue position of the pearl allele mutation is based on this re-annotation.

### Expression analysis from RNA-seq data

RNA-sequencing data for whole embryos and adult tissues (retina, liver, olfactory epithelium) were obtained from previously described datasets deposited in the SRA database with sequence accessions SRR5878849-SRR5878856 (Holt et al. 2018). For HH25 Oriental Frill and Racing Homer embryo heads, RNA from whole embryonic heads was isolated using the Qiagen RNEasy Kit, and submitted to the University of Utah High Throughput Genomics Shared Resource for Illumina TruSeq stranded library preparation. Libraries were sequenced on the Illumina HiSeq platform. Data are available in NCBI Sequence Read Archive under BioProject PRJNA680754

We mapped reads to the Cliv_2.1 reference genome using STAR (Dobin et al. 2013), and counted reads in features using FeatureCounts (Liao et al. 2014). Reads per million for the *SLC2A11B* gene were calculated based on total number of uniquely mapped reads per sample. For each HH25 embryo head, we looked at reads overlapping two SNPs within the pearl eye haplotype (ScoHet5_1307:1895834 and ScoHet5_1307:1896042) to predict genotypes. We evaluated relative expression level of *SLC2A11B* between orange and pearl alleles using a two-tailed T-test to compare reads per million in each sample set.

### Protein conservation, structure prediction, and mutation evaluation

We obtained protein sequences for SLC2A11B orthologues across species using NCBI blastp and generated multi-species alignments using Clustal Omega (Madeira et al. 2019), and then visualized using Jalview2 (Waterhouse et al. 2009). We assessed the predicted structure of the SLC2A11B protein by using Phobius (Käll et al. 2004) to predict cytoplasmic, non-cytoplasmic, transmembrane, and signal peptide domains. As the premature stop codon in *SLC2A11B* occurs very early in the protein sequence, we evaluated the likely impact of the premature stop codon by identifying the next in-frame methionine where translation could initiate to make the longest possible partial protein. We input this truncation into PROVEAN (Choi et al. 2012; Choi and Chan 2015) as a deletion of the first 95 amino acids.

### Gene ontology analysis

We mapped gene ontology annotations to identifiers for genes within the two bull eye candidate regions using DAVID (https://david.ncifcrf.gov/; Huang et al. 2009). We used annotations from Biological Process (GOTERM_BP_ALL; GOTERM_BP_DIRECT), Cellular Component (GOTERM_CC_ALL; GOTERM_CC_DIRECT), and Molecular Function (GOTERM_MF_ALL; GOTERM_MF_DIRECT) gene ontology databases, and searched results for GO terms containing the keywords “pigment”, “melanosome” or “melanocyte”.

## Data Availability

Whole genome sequencing and RNA-sequencing datasets generated for this study have been deposited to the NCBI SRA database under BioProject PRJNA680754. Previously generated whole genome sequencing and RNA-seq data used in this study is available under BioProject accessions PRJNA513877, PRJNA428271, and PRJNA167554.

## Acknowledgements and Funding Information

We thank current and former members of the Shapiro lab for assistance with sample collection and processing. We thank Eric Domyan, Anna Vickrey, Hannah Van Hollebeke, Alexa Davis, Tennyson George, Marissa Burton, and Lucas Periera for technical assistance and advice. Layne Gardner generously shared the Pomeranian Pouter and Scandaroon photographs featured in Fig. 3A. We thank members of the Utah Pigeon Club for providing samples. We acknowledge a computer time allocation from the University of Utah Center for High Performance Computing. This work was supported by the National Institutes of Health (R35GM131787 to M.D.S. and F32DE028179 to E.B.), the National Science Foundation (GRF 1256065 to R.B.), the Jane Coffin Childs Memorial Fund for Medical Research (fellowship to E.M.), and the University of Utah Undergraduate Research Opportunities Program (fellowship support to B.P., R.W., and T.G.).

