## Supplemental Figures 1-4 for "Two Genomic Loci Control Three Eye Colors in the Domestic Pigeon (*Columba livia*)"

**Affiliations:**
[1]School of Biological Sciences, University of Utah, Salt Lake City, UT 84112, USA
[2] Department of Human Genetics and USTAR Center for Genetic Discovery, University of Utah, Salt Lake City, UT, USA

*Author for Correspondence:
Michael D. Shapiro, School of Biological Sciences, 257 South 1400 East, Salt Lake City, UT 84112 USA; phone +1 801 581 5690;

Figure S1
Figure S2
Figure S3
Figure S4

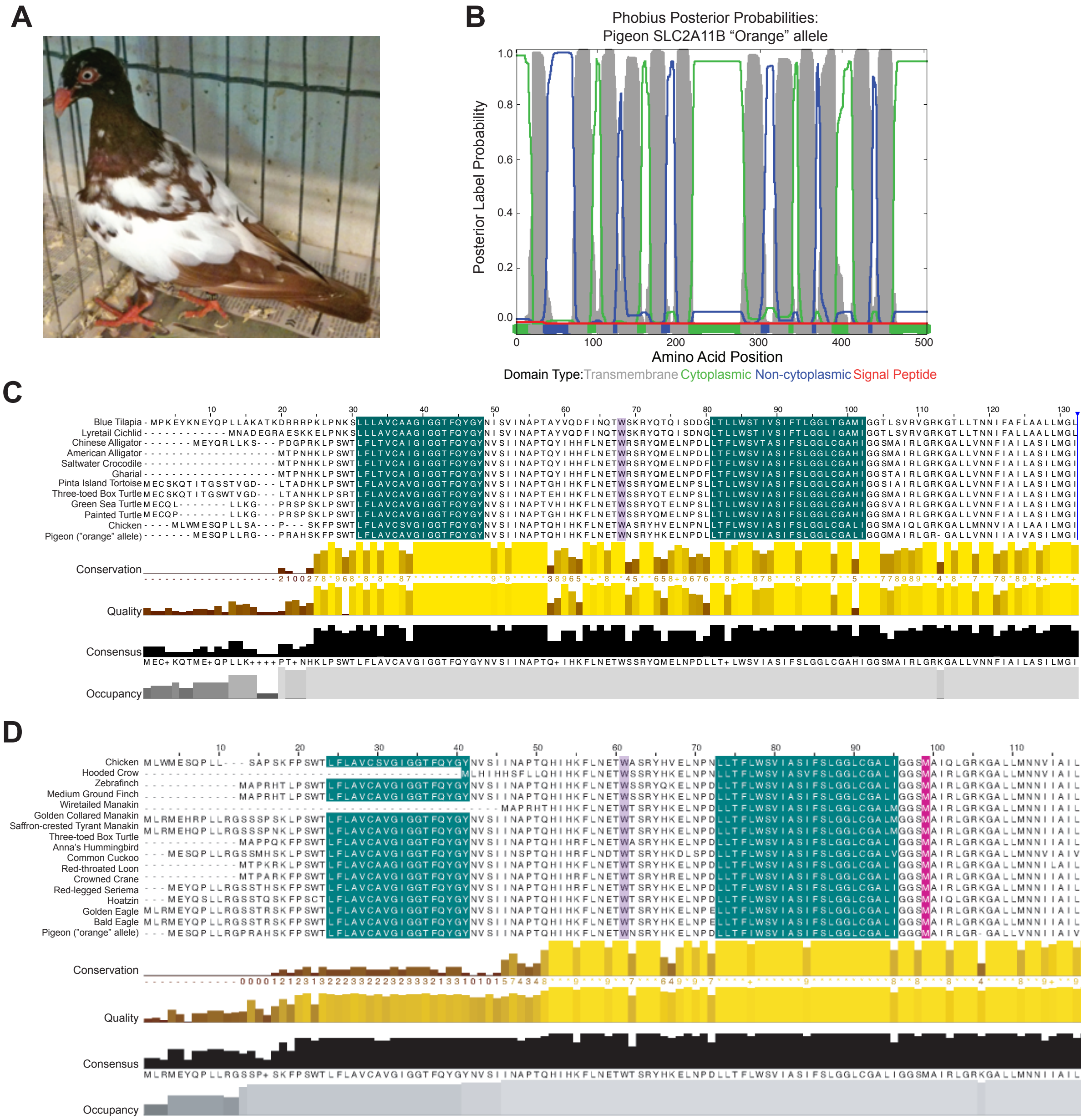


**Figure S1. A premature stop codon in exon 3 of SLC2A11B is predicted to disrupt conserved protein structure and function.** (A) The pigeon sequenced for the *C. livia* reference genome, a Danish Tumbler with pearl eyes [from Shapiro et al. 2013]. (B) Sequence-based domain prediction of SLC2A11B structure from Phobius. Amino acid positions of protein regions are depicted on the X axis and are color-coded by predicted region type. Probability scores for each predicted domain type are shown on the Y axis. (B) Multi-species alignment of SLC2A11B protein sequence across 12 fish and sauropsid species, including pigeon. Teal color indicates the spans of the first two transmembrane domains predicted by Phobius, which are highly conserved across species. The residue affected by the pearl eye mutation, W58 in the pigeon orange allele protein sequence, is highlighted in purple. (D) Multi-species alignment of SLC2A11B protein sequence across 17 bird species, including pigeon. Teal and purple indicate same features as in (C). Pink indicates the next in-frame methionine after the pearl eye W58X mutation from which translation could initiate.

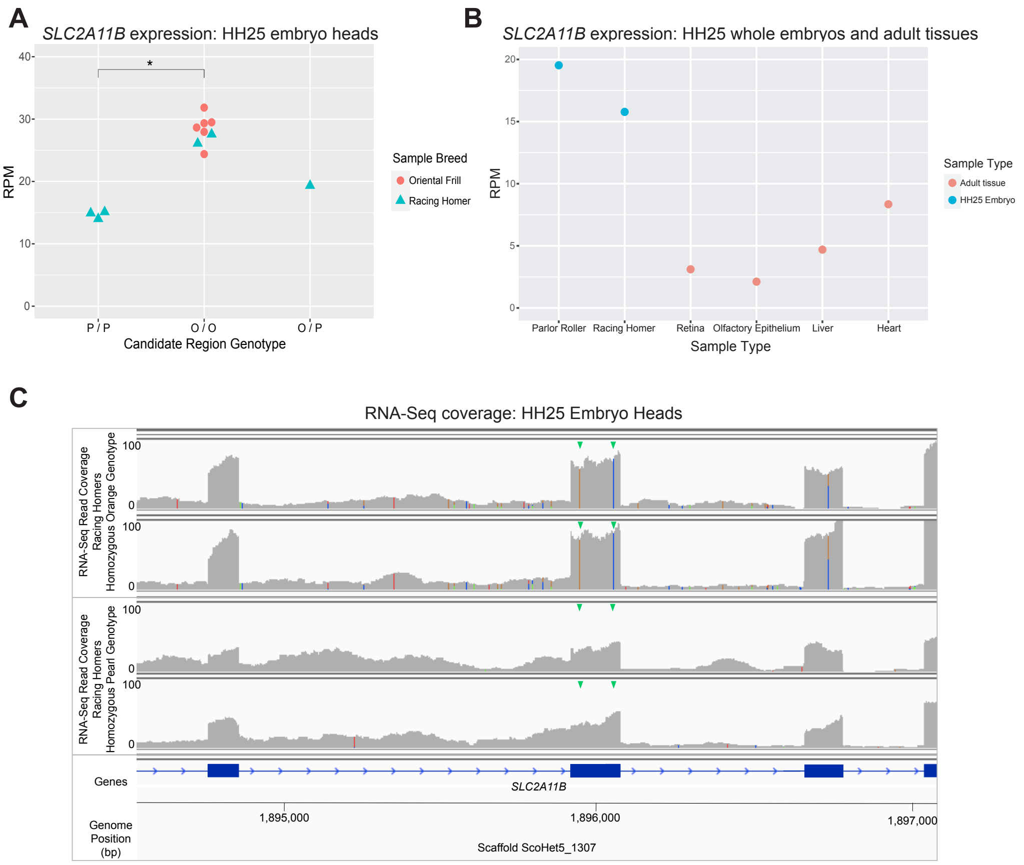


**Figure S2. RNA-Seq shows a reduction in spliced *SLC2A11B* transcripts in embryos carrying the Pearl allele.** (A) Plot of normalized *SLC2A11B* read count in RNA-sequencing data from whole HH25 embryo heads of two different breeds, Racing Homer and Oriental Frill. Y axis shows normalized read count, calculated as *SLC2A11B* reads per million uniquely mapped RNA-seq reads. Expression is significantly reduced (p = 1.139e-07, two-tailed t test) in embryos homozygous for the pearl allele (P/P, n=3) compared to embryos homozygous for the orange allele (O/O, n=8). One heterozygous sample (O/P) shows an intermediate phenotype. (B) Plot of normalized *SLC2A11B* read count in RNA-sequencing data from whole HH25 embryos (one Parlor Roller, one Racing Homer) and four adult tissues (data from Holt et al. 2018). Y axis shows normalized read count, calculated as *SLC2A11B* reads per million uniquely mapped RNA-seq reads (C) Visual evaluation of mapped RNA-seq reads shows an apparent increase in the proportion of intronic reads in samples homozygous for the pearl allele. Each track represents a single sample. Y axis shows non-normalized RNA-seq coverage. X axis shows basepair positions across the region, *SLC2A11B* exons are indicated in blue. Green arrowheads point to the locations of two exonic single nucleotide polymorphisms associated with eye color. These SNPs are highlighted in the orange genotype samples (upper two tracks) as the sequence in these samples differs from the pearl-eyed reference genome. Additional colored lines in each sample mark non-reference alleles.

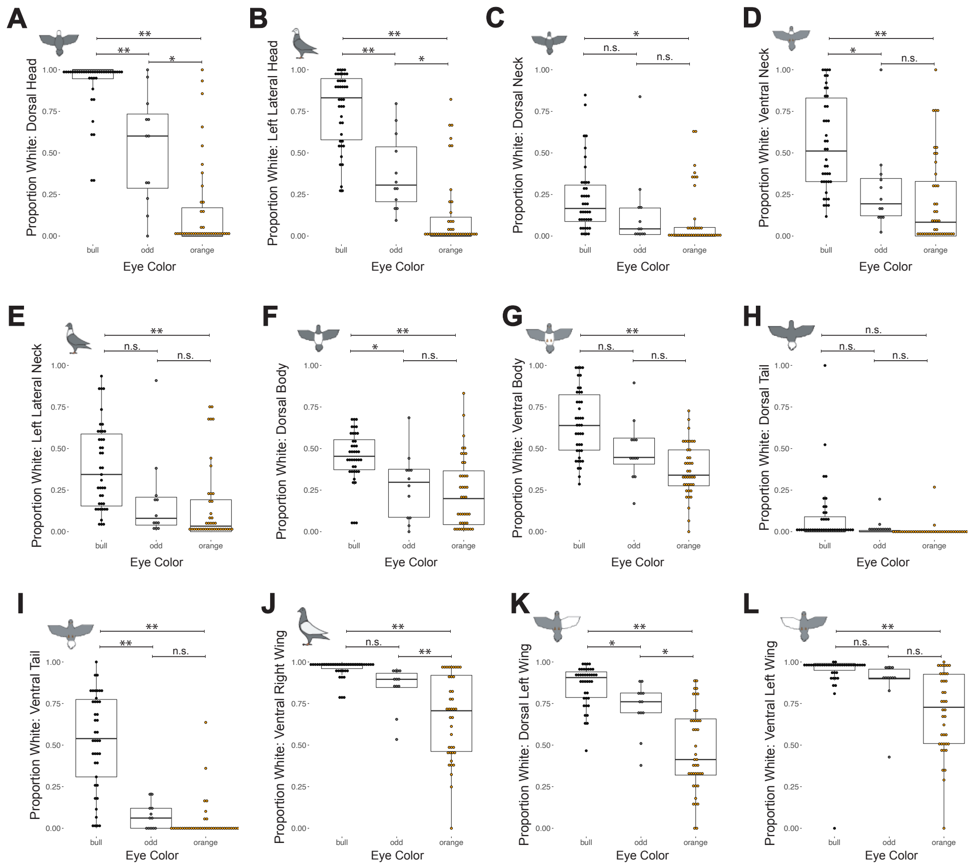


**Figure S3. Association between plumage pigmentation and eye color in Pomeranian Pouter x Scandaroon $F_2$ offspring.** (A-L) Boxplots depicting the proportion of white plumage on the indicated body region in Pomeranian Pouter x Scandaroon $F_2$ birds with bull, odd, or orange eyes. **, $p \leq 0.0001$; *, $0.001 < p \leq 0.01$; n.s., $p > 0.01$. Boxes span from the first to third quartile of each data set, with lines indicating the median. Whiskers span up to 1.5x the interquartile range.

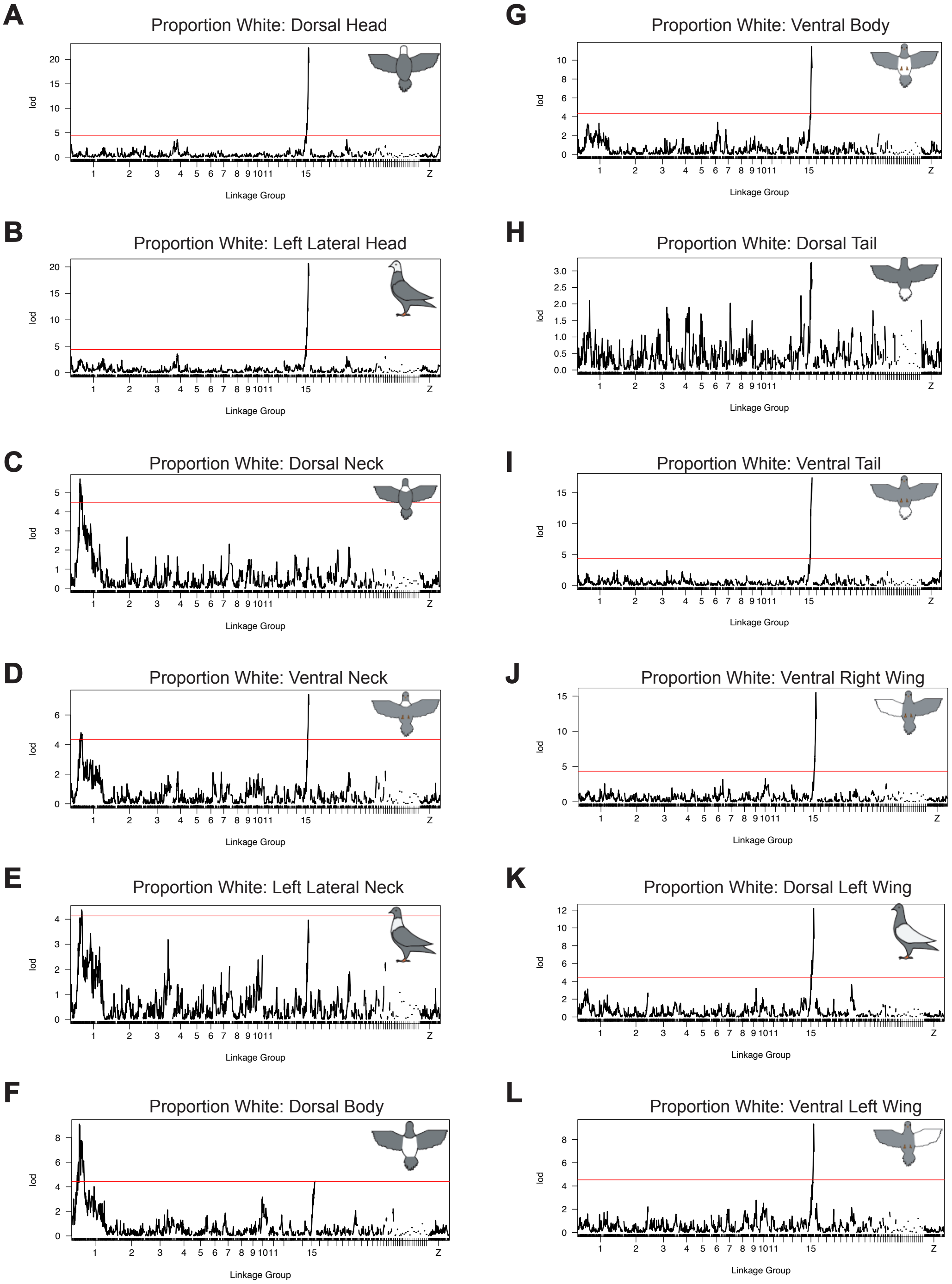


**Figure S4. Two linkage groups are associated with regionally-specific white plumage in a Pomeranian Pouter x Scandaroon $F_2$ intercross.** (A-L) Genome-wide QTL scans for proportion of white plumage on the indicated body region. Red lines indicate the 5% genome-wide significance threshold. In (H), a significant QTL was not identified for the dorsal tail region.
